# Control of bacterial immune signaling by a WYL domain transcription factor

**DOI:** 10.1101/2022.01.04.474952

**Authors:** Chelsea L. Blankenchip, Justin V. Nguyen, Rebecca K. Lau, Qiaozhen Ye, Yajie Gu, Kevin D. Corbett

## Abstract

Bacteria use diverse immune systems to defend themselves from ubiquitous viruses termed bacteriophages (phages). Many anti-phage systems function by abortive infection to kill a phage-infected cell, raising the question of how they are regulated to avoid cell killing outside the context of infection. Here, we identify a transcription factor associated with the widespread CBASS bacterial immune system, that we term CapW. CapW forms a homodimer and binds a palindromic DNA sequence in the CBASS promoter region. Two crystal structures of CapW suggest that the protein switches from an unliganded, DNA binding-competent state to a ligand-bound state unable to bind DNA. We show that CapW strongly represses CBASS gene expression in uninfected cells, and that phage infection causes increased CBASS expression in a CapW-dependent manner. Unexpectedly, this CapW-dependent increase in CBASS expression is not required for robust anti-phage activity, suggesting that CapW may mediate CBASS activation and cell death in response to a signal other than phage infection. Our results parallel concurrent reports on the structure and activity of BrxR, a transcription factor associated with the BREX anti-phage system, suggesting that CapW and BrxR are members of a family of universal defense signaling proteins.

## Introduction

The constant evolutionary arms race between bacteria and the viruses that infect them, called bacteriophages (phages), has resulted in the evolution of a broad array of bacterial immune systems. These include the well-characterized restriction-modification and CRISPR/Cas systems, but also less well-understood systems like BREX (<u>B</u>acte<u>r</u>iophage <u>Ex</u>clusion) (1,2), CBASS (<u>C</u>yclic Oligonucleotide-<u>B</u>ased <u>A</u>nti-Phage <u>S</u>ignaling <u>S</u>ystem) (3,4), and others (5,6). Many bacterial immune systems, including CBASS, act by a so-called abortive infection mechanism and kill an infected cell to avoid phage reproduction (3,7). Because of their destructive power, abortive infection systems must be exquisitely tuned to avoid activation in an uninfected cell, but activate rapidly and reliably upon infection.

CBASS immune systems are widespread and extremely diverse, with over 6,200 distinct systems identified to date across bacteria and archaea (3,8). CBASS systems show diversity in both their activation mechanisms and their cell-killing mechanisms. All CBASS systems encode an oligonucleotide cyclase related to the mammalian innate-immune sensor cGAS (cyclic GMP-AMP synthase) that synthesizes a cyclic di- or trinucleotide second messenger molecule (8,9), and a second-messenger activated effector protein that mediates cell death to abort the viral infection. Type I CBASS systems encode only these two proteins, suggesting that their cGAS-like enzyme has an innate infection-sensing capability (10). The recent discovery of Type I CBASS systems that encode a cGAS-like enzyme and an effector related to eukaryotic STING (<u>ST</u>imulator of <u>In</u>terferon <u>G</u>enes) suggests that the mammalian cGAS-STING innate-immune pathway evolved from these systems (11).

The majority of bacterial CBASS systems encode putative regulators that likely provide an additional level of control over their activation. Type II CBASS systems encode enzymes related to eukaryotic ubiquitin-transfer machinery, whose roles in signaling remain unknown (4,10). In Type III CBASS systems, peptide-binding HORMA domain proteins and a AAA+ATPase, Trip13, regulate cGAS activation through HORMA-peptide binding (7). These regulators are thought to represent the evolutionary precursors of the diverse HORMA domain signaling protein family in eukaryotes, which regulate key cell-cycle checkpoints, DNA repair and recombination in mitotic and meiotic cells, and autophagy signaling (7,12).

The diversity of bacterial CBASS systems suggests that additional mechanisms of phage infection sensing and signaling regulation in these systems remain to be discovered. Here, we identify a novel transcription factor, CapW, that is associated with hundreds of distinct CBASS systems. We show that CapW is a transcriptional repressor that binds the promoter region of its cognate CBASS operon to inhibit expression of CBASS genes. Two structures of CapW from different bacteria reveal a dimeric assembly that likely binds a small-molecule ligand to control DNA binding and transcriptional repression. Structure-based mutagenesis of CapW reveals that the protein’s putative ligand-binding WYL domain is required for a phage infection-dependent increase in CBASS expression, but that this increase is unexpectedly not required for robust anti-phage activity. These data suggest that CapW may mediate increased CBASS expression, and concomitant host-cell toxicity, in response to a signal other than phage infection. Parallel discovery of CapW-like transcription factors in other bacterial immune systems including BREX (13,14) suggests that CapW and its relatives make up a family of ligand-responsive transcriptional switches, which directly sense stress signals and activate expression of diverse immune systems.

## Materials and Methods

### Bioinformatics

To comprehensively search CBASS systems for homologs of *E. coli* upec-117 CapW (NCBI #WP_001534693.1), we exported the genomic DNA sequence +/-10 kb of 6233 previously-reported CD-NTases (3) using the Integrated Microbial Genomes (IMG) database at the DOE Joint Genome Institute (https://img.jgi.doe.gov). We used NCBI Genome Workbench (https://www.ncbi.nlm.nih.gov/tools/gbench/) to perform custom TBLASTN searches for proteins related to *E. coli* upec-117 CapW. CBASS system type and effector assignments for each hit were taken from Cohen et al. (3) and manually updated. Each hit was manually inspected for the presence of CapW specifically associated with CBASS rather than a neighboring operon.

### Protein Expression and Purification

Codon-optimized sequences encoding full-length CapW from *E. coli* upec-117 (NCBI #WP_001534693.1), *P. aeruginosa* PA17 (NCBI #WP_023098969.1), and S. *maltophilia* C11 (IMG #2657474953) were synthesized (Invitrogen/GeneArt) and cloned into UC Berkeley Macrolab vector 2BT (Addgene #29666) to generate constructs with N-terminal TEV protease-cleavable His_6_-tags. Point mutations were generated by PCR-based mutagenesis.

Proteins were expressed in *E. coli* strain Rosetta 2 (DE3) pLysS (EMD Millipore). Cultures were grown at 37°C to A_600_=0.7, then induced with 0.25 mM IPTG and shifted to 20°C for 16 hours. Cells were harvested by centrifugation and resuspended in buffer A (25 mM Tris pH *7.5*, 10% glycerol, and 1 mM NaN_3_) plus 300 mM NaCl, 5 mM imidazole, 5 mM ß-mercaptoethanol. Proteins were purified by Nibaffinity (Ni-NTA agarose, Qiagen) then passed over an anion-exchange column (Hitrap Q HP, GE Life Sciences) in Buffer A plus 100 mM NaCl and 5 mM β-mercaptoethanol, collecting flow-through fractions. Tags were cleaved with TEV protease (15), and cleaved protein was passed over another Ni^2+^ column (collecting flow-through fractions) to remove uncleaved protein, cleaved tags, and tagged TEV protease. The protein was passed over a size exclusion column (Superdex 200, GE Life Sciences) in buffer GF (buffer A plus 300 mM NaCl and 1 mM dithiothreitol (DTT)), then concentrated by ultrafiltration (Amicon Ultra, EMD Millipore) to 10 mg/ml and stored at 4°C. All point mutants showed equivalent migration on size exclusion column compared to wild type. For selenomethionine derivatization, protein expression was carried out in M9 minimal media supplemented with amino acids plus selenomethionine prior to IPTG induction (16).

For characterization of oligomeric state by size exclusion chromatography coupled to multiangle light scattering (SEC-MALS), 100 μL of purified *P. aeruginosa* PA17 CapW, S. *maltophilia* C11 CapW, or *E. coli* upec-117 CapW at 5 mg/mL was injected onto a Superdex 200 Increase 10/300 GL column (GE Life Sciences) in buffer GF. Light scattering and refractive index profiles were collected by miniDAWN TREOS and Optilab T-rEX detectors (Wyatt Technology), respectively, and molecular weight was calculated using ASTRA v. 6 software (Wyatt Technology).

### Crystallization and Structure Determination

For crystallization of *P. aeruginosa* PA17 CapW, selenomethionine-derivatized protein in buffer GF (9 mg/mL) was mixed 1:1 with well solution containing 0.27 M LiSO_4_, 1% PEG 400, and 0.1 M sodium acetate (pH 5.0) in hanging-drop format. Crystals were cryoprotected by the addition of 30% glycerol, and flash-frozen in liquid nitrogen. Diffraction data were collected at the Advanced Photon Source NE-CAT beamline 24ID-E (see support statement below) and processed with the RAPD data-processing pipeline (https://github.com/RAPD/RAPD), which uses XDS (17) for data indexing and reduction, AIMLESS (18) for scaling, and TRUNCATE (19) for conversion to structure factors. We determined the structure by single-wavelength anomalous diffraction methods in the PHENIX Autosol wizard (20). We manually rebuilt the initial model in COOT (21), and refined in phenix.refine (22) using positional and individual B-factor refinement **(Table S3).**

For crystallization of S. *maltophilia* C11 CapW, protein in a buffer containing 20mM Tris pH 8.5, 1 mM DTT, and 100 mM NaCl (14mg/mL) was mixed 2:1 with well solution containing 0.1 M Tris pH 8.5, and 1.5 M Lithium sulfate in sitting drop format. Crystals were cryoprotected by the addition of 24% glycerol and flash-frozen in liquid nitrogen. Diffraction data were collected at the Advanced Light Source BCSB beamline 5.0.2 (see support statement below) and processed with the DIALS data-processing pipeline (https://dials.github.io) (23). We determined the structure by molecular replacement in PHASER (24), using individual wHTH and WYL domain structures from *P. aeruginosa* PA17 CapW as search models. We manually rebuilt the initial model in COOT (21), and refined in phenix.refine (22) using positional and individual B-factor refinement **(Table S3).**

### DNA binding

For electromobility shift DNA-binding assays (EMSA), the S. *maltophilia* C11 CBASS promoter region was amplified via PCR with one primer 5’-labeled with 6-carboxyfluorescein (5’-6-FAM), followed by gel-purification (Machery-Nagel Nucleospin). For the EMSA control, a random sequence from within the S. *maltophilia* C11 CapW gene that was similar in length and GC content to the S. *maltophilia* C11 promoter region was amplified via PCR with one primer 5’-labeled with 6-carboxyfluorescein (5’-6-FAM), followed by gel-purification (Machery-Nagel Nucleospin). 10 μL reactions with 100 nM DNA and indicated concentrations of protein were prepared in a buffer containing 50 mM Tris-HCl pH 8.5, 50 mM NaCl, 5 mM MgCl_2_, 5% glycerol, and 1 mM DTT. After a 1 hour incubation at room temperature, reactions were loaded onto 2% TBE-agarose gels in running buffer with 0.5X TBE pH 8.5 running buffer, run for 2 hours at 60 V at 4°C, and imaged using a Bio-Rad ChemiDoc system (Cy2 filter settings).

For DNA binding fluorescence polarization (FP) assays, a 30 bp double-stranded DNA was produced by annealing complementary oligos, one of which was 5’-6-FAM labeled. Binding reactions (30 μL) contained 25 mM Tris pH 8.5, 50 mM NaCl, 5% glycerol, 5 mM MgCl_2_, 1 mM DTT, 0.01% nonidet p40 substitute, 50 nM DNA, and the indicated amounts of protein. After a 10 minute incubation at room temperature, fluorescence polarization was read using a Tecan Infinite M1000 PRO fluorescence plate reader, and binding data were analyzed with Graphpad Prism v.9.2.0 using a single-site binding model.

### GFP Reporter Assays

To generate a GFP reporter plasmid, a DNA sequence encoding full-length *E. coli* upec-117 CapW and its promoter, adjacent to a gene encoding super-folder GFP was synthesized (IDT) and cloned via isothermal assembly into pBR322. Point mutations were generated by PCR-based mutagenesis. Vectors were transformed into *E. coli* strain JP313 (25). 100 μL of saturated overnight culture was added to 5 mL LB broth plus ampicillin and grown at 37°C to an OD_600_ of 0.4-0.5. 500 μL of culture was pelleted by centrifugation and resuspended in 100 μL of 2x SDS-PAGE loading buffer (125 mM Tris pH 6.8, 20% Glycerol, 4% SDS, 200 mM DTT, 180 μM bromophenol blue). Samples were boiled for 5 minutes, then 1-10 μL was loaded onto an SDS-PAGE gel. Proteins were transferred to a PVDF membrane (Bio-Rad Trans-Blot Turbo), then the membranes were blocked with 5% nonfat dry milk and blotted for appropriate proteins. Blots were imaged using a Bio-Rad ChemiDoc system using filters to image horseradish peroxidase activity. Antibodies used: Mouse anti-GFP primary antibody (Roche) at 1:3,000 dilution; Mouse anti-FLAG primary antibody (Sigma-Aldrich) at 1:3,000 dilution; Mouse anti-RNA polymerase primary antibody (clone NT63; BioLegend #10019-878) at 1:3000 dilution; Goat anti-mouse HRP-linked secondary antibody (Millipore Sigma) at 1:30,000 dilution.

To measure reporter responses to phage infection, GFP reporter plasmids were transformed into *E. coli* strain JP313 (25). Overnight cultures were diluted into fresh LB media with ampicillin and grown to an OD_600_ of 0.25-0.35. Bacterial cultures were infection with bacteriophage λ cI-diluted in phage buffer plus 1 mM CaCl_2_ at a multiplicity of infection of 10. Cultures were incubated at 30°C and timepoints were taken immediately after infection (0 minutes), and 30, 60, 90, and 120 minutes post infection. Samples were prepared and analyzed by Western blot as above.

### Bacteriophage infection assays

To measure anti-phage activity of *E. coli* upec-117 CBASS, we transformed *E. coli* strain JP313 (25) with a pLOCO2 plasmid encoding the entire CBASS system from this species (bases 69873 to 76528 of NCBI accession # CP054230.1, encoding proteins QKY44554.1 (Cap17;MTA/SAH nucleoside phosphorylase), QKY44555.1 (CapW), QKY44556.1 (CdnC), QKY44557.1 (Cap7;HORMA), QKY445558.1 (Cap6; TRIP13), QKY44559.1 (Cap18; 3’-5’ exonuclease), and QKY44560.1 (Cap19; 3-transmembrane protein) (gift from A. Whiteley) or the mutants noted in **Figure 5d-e.** As a negative control, we used the same pLOCO2 backbone encoding lacI and sfGFP. For bacteriophage infection plaque assays, a single bacterial colony or equivalent mass was picked from a fresh LB agar plate and grown at 37°C in LB broth plus ampicillin to an OD_600_ of 0.2-0.3. 500 μL of cells were mixed with 4.5 mL of 0.35% LB top agar with carbenicillin, then poured onto LB plates containing carbenicillin. Plates were spotted with 3 μL of bacteriophage λcl-(cI gene deleted to inhibit lysogeny and promote clear plaque formation) diluted in phage buffer (150 mM NaCl, 40 mM Tris pH 7.5, 10 mM MgSO_4_) plus 1 mM CaCl_2_ at 6 dilutions: ~4×10^7^ PFU/mL and 5 10-fold dilutions thereof. Plates were incubated at 37°C for 18 hours, then imaged.

**Figure 1.**
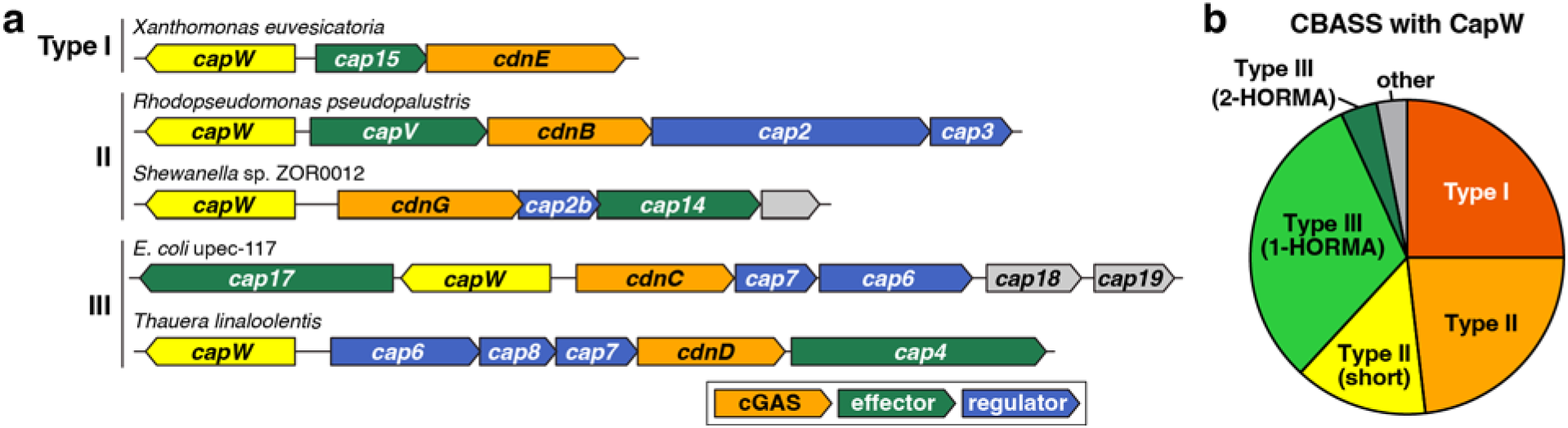
Identification of CapW in CBASS systems. (a) Operon schematics of five CBASS systems encoding *capW* genes (yellow). Each system’s CBASS type is noted at left, and genes are color-coded with cGAS-like enzymes orange, effectors green, regulators blue, and proteins with unknown function gray. cGAS-like enzymes are labeled according to their clade, as previously assigned (8). See **Table S1** for a comprehensive list of CapW-associated CBASS systems, and **Table S2** for gene names and descriptions. (b) Pie chart showing the number of CapW-associated CBASS systems, with Type I red, Type II orange, Type II (short) yellow, Type III (1-HORMA) light green, Type III (2-HORMA) dark green, and other/unknown gray (see **Table S1).**

**Figure 2.**
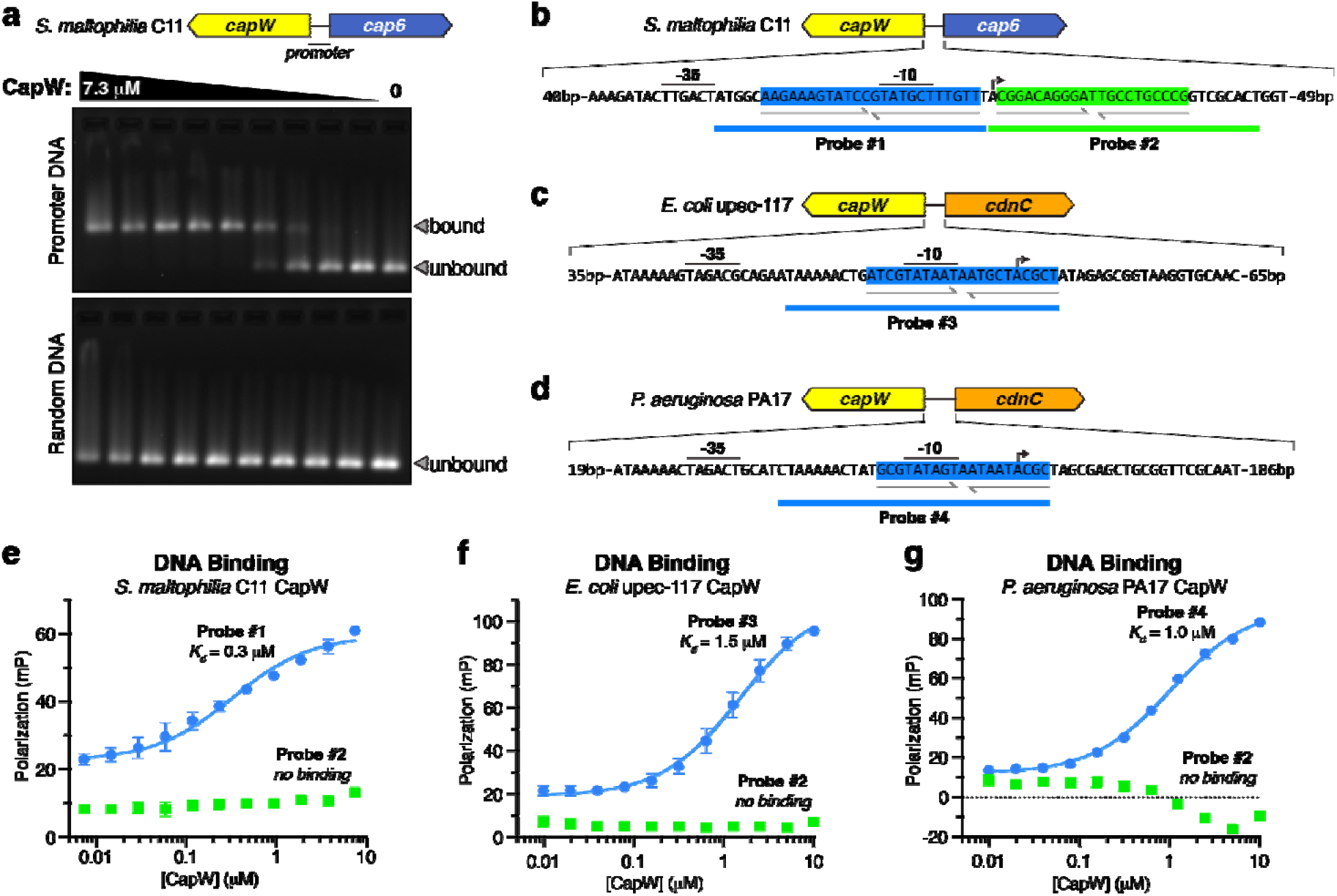
CapW binds the CBASS promoter region. (a) Electrophoretic mobility shift assay (EMSA) for *Sm* CapW binding the full promoter region of its cognate CBASS system (upper panel), or an equivalent-length random DNA (lower panel). The leftmost lane contains 7.3 μM CapW (monomer), and lanes 2-9 contain progressive 2-fold dilutions. Estimated *Kd* = 0.2-0.3 μM. (b) Diagram of the S. *maltophilia* C11 CBASS promoter region, with two palindromic sequences highlighted in blue and green. Two 30-bp DNA probes used for fluorescence polarization binding assays are also shown. Promoter sequences (−35, −10, and TSS sites) were predicted with the BPROM server (40). (c) Diagram of the *E. coli* upec-117 CBASS promoter region, with palindromic sequence highlighted in blue. (d) Diagram of the *P. aeruginosa* PA17 CBASS promoter region, with palindromic sequence highlighted in blue. (e) Fluorescence polarization assay showing robust binding of *Sm* CapW to Probe #1 (blue, *Kd* = 0.3 +/− 0.1 μM), but not Probe #2 (green). For panels (e), (f), and (g), datapoints represent the mean +/− standard deviation of three measurements. (f) Fluorescence polarization assay showing binding of *Ec* CapW to Probe #3 (blue: *Kd* = 1.5 +/− 0.3 μM), but not Probe #2 from S. *maltophilia* C11 CBASS (green). (g) Fluorescence polarization assay showing binding of *Pa* CapW to Probe #4 (blue: *Kd* = 1.0 +/− 0.1 μM), but not Probe #2 from *S. maltophilia* C11 CBASS (green).

**Figure 3.**
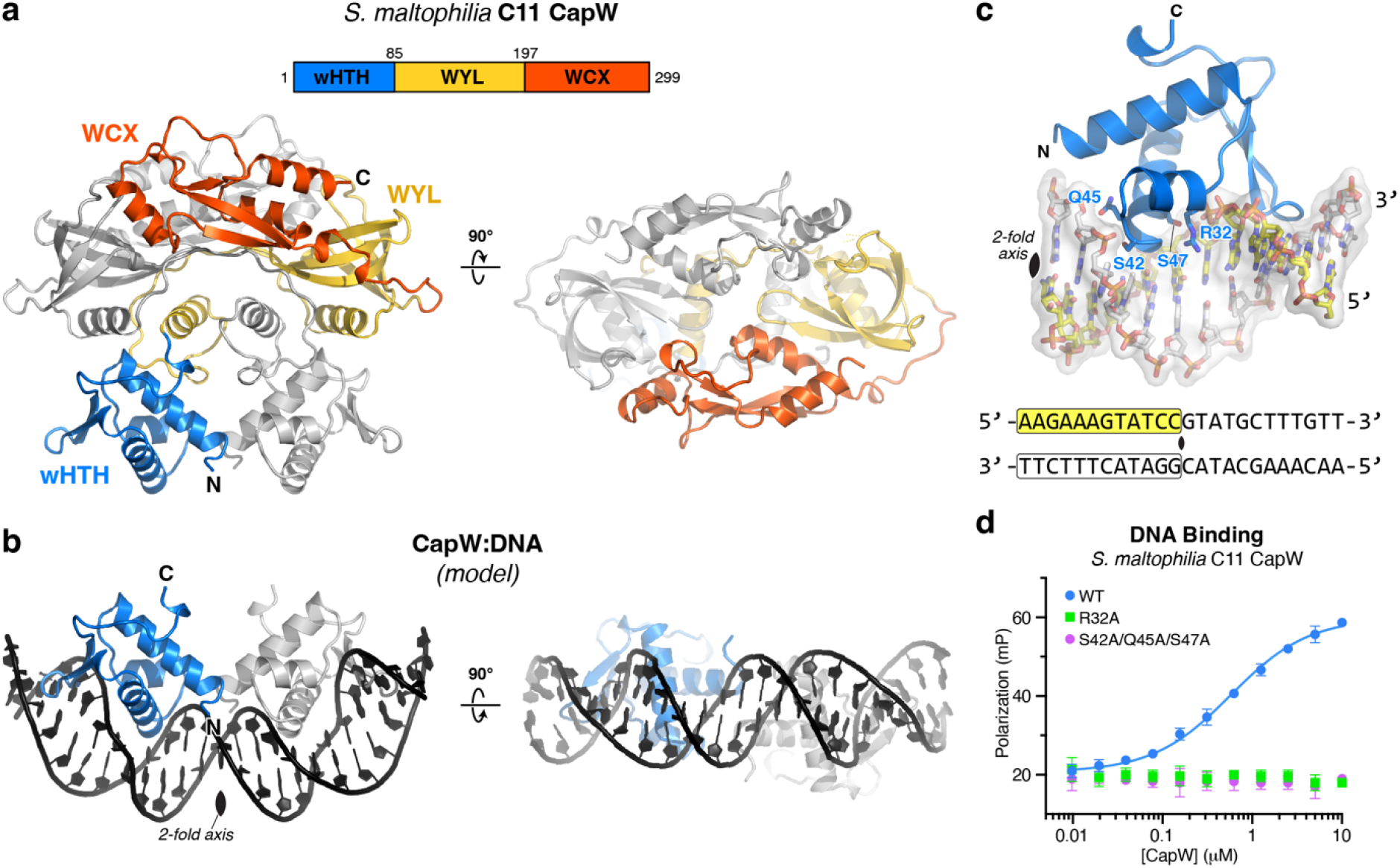
Structure of *Sm* CapW. (a) Structure of the *Sm* CapW dimer, with wHTH domain colored blue, WYL domain yellow, and WYL C-terminal extension (WCX) domain orange. (b) Model of the CapW wHTH domains binding DNA, generated by docking the structure of DNA-bound *Acinetobacter* BrxR (PDB ID 7T8K) (13) onto the structure of *Sm* CapW (overall Cα r.m.s.d. 2.9 Å; see **Fig. S2d).** (c) Closeup view of the *Sm* CapW wHTH domain-DNA model, with residues chosen for mutagenesis labeled. The DNA sequence modeled is one half of the site identified in **Fig. 1b,** and the two-fold rotation axis of the overall complex and DNA is marked with a black oval. (d) Fluorescence polarization assay showing binding of Sm CapW (WT and indicated mutants) to Probe #1 from **Fig. 2b.**

**Figure 4.**
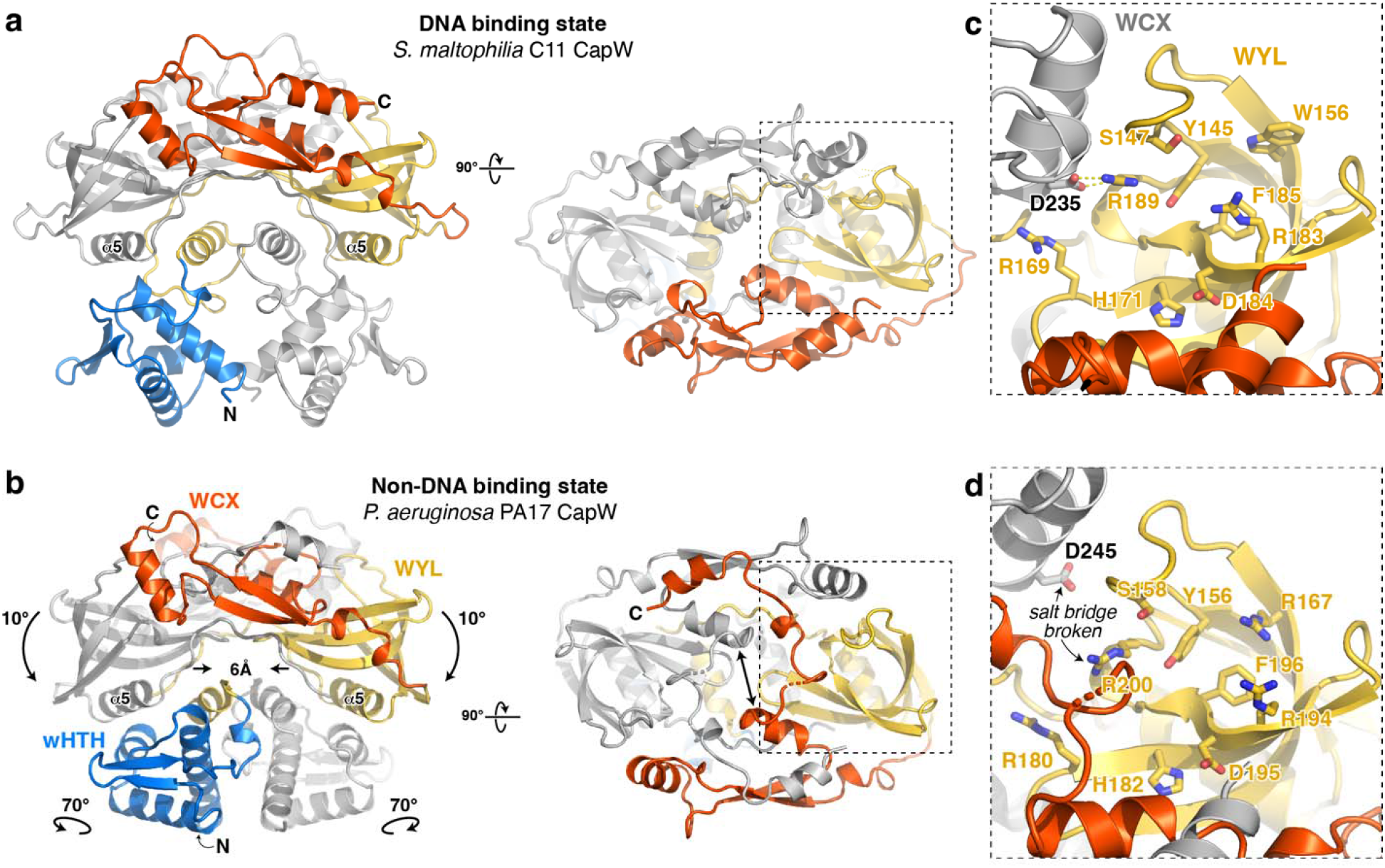
The structure of *Pa* CapW reveals a non-DNA binding state. (a) Structure of Sm CapW, with wHTH domain colored blue, WYL domain yellow, and WYL C-terminal extension (WCX) domain orange. (b) Low-pH structure of *Pa* CapW, with domains colored as in panel (a). (c) Closeup of the putative ligand-binding site on the Sm CapW WYL domain. WYL domain residues are labeled in yellow, and residues from the dimer-related WCX domain (gray) are labeled in black. See **Fig. S3** for surface conservation. (d) Closeup of the putative ligand-binding site on the *Pa* CapW WYL domain. WYL domain residues are labeled in yellow, and residues from the dimer-related WCX domain (gray) are labeled in black. The arginine-asparate salt bridge observed between WYL and the dimer-related WCX domain in Sm CapW is broken in this structure (arrow).

**Figure 5.**
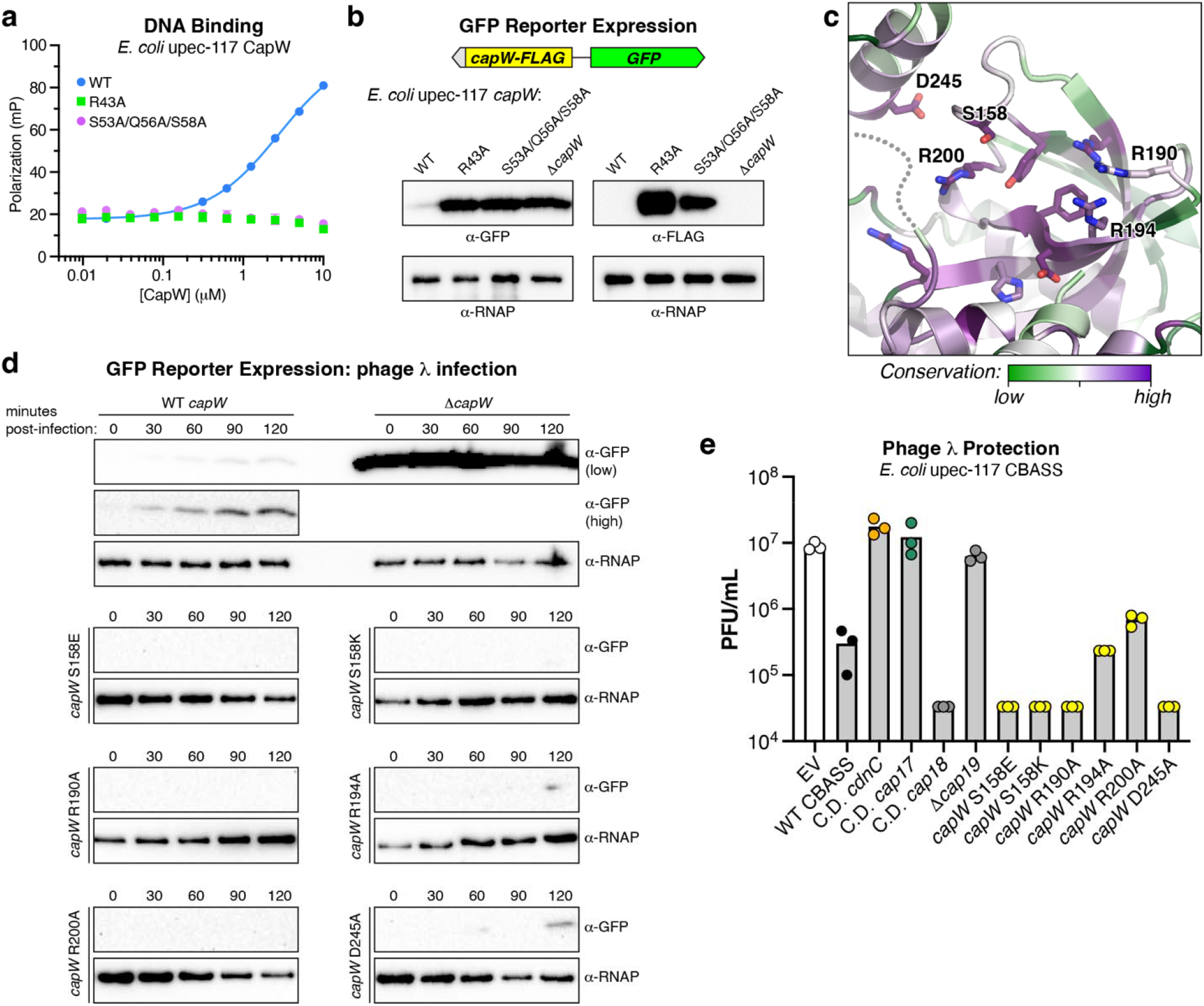
CapW is a transcriptional repressor. (a) Fluorescence polarization assay showing binding of *Ec* CapW (WT and indicated mutants) to Probe #3 from **Fig. 2c.** (b) Reporter assay with *E. coli* upec-117 CapW and promoter region, with GFP in place of the CBASS core genes. Western blots show expression of GFP (left) and CapW-FLAG (right) in log-phase cells in the presence of wild-type or mutant CapW. Δ*capW:* double-stop codon inserted after codon 10; α-RNAP: anti-RNA polymerase western blot loading control. See **Fig. S5** for a diagram of the *E. coli* upec-117 CBASS promoter with both forward and reverse promoter sequences. (c) Top-down view of the *Pa* CapW WYL domain, colored by conservation (see **Fig. S3c-d).** Highly conserved putative ligandinteracting residues mutated in *Ec* CapW are labeled (residue numbering is equivalent for *Pa* CapW and *Ec* CapW in this region). (d) Reporter assay showing expression changes in λ cI-infected cells (multiplicity of infection=10) at the indicated times post-infection. (e) Quantification of plaque assays showing infectivity of λ cI-against the *E. coli* upec-117 CBASS system and indicated mutants, expressed as plaque-forming units (PFU) per mL of phage stock. Bars represent the average of three measurements (colored circles). Catalytic-dead (C.D.) *cdnC:* D73N/D75N; C.D. *cap17:* D472N; C.D. *cap18:* D165N; Δ*cap19* (double-stop codon inserted after codon 10). See **Fig. S6** for full data.

### APS NE-CAT Support Statement

This work is based upon research conducted at the Northeastern Collaborative Access Team beamlines, which are funded by the National Institute of General Medical Sciences from the National Institutes of Health (P30 GM124165). The Eiger 16M detector on the 24-ID-E beam line is funded by a NIH-ORIP HEI grant (S10OD021527). This research used resources of the Advanced Photon Source, a U.S. Department of Energy (DOE) Office of Science User Facility operated for the DOE Office of Science by Argonne National Laboratory under Contract No. DE-AC02-06CH11357.

### ALS BCSB Support Statement

The Berkeley Center for Structural Biology is supported in part by the Howard Hughes Medical Institute. The Advanced Light Source is a Department of Energy Office of Science User Facility under Contract No. DE-AC02-05CH11231. The Pilatus detector on 5.0.1. was funded under NIH grant S10OD021832. The ALS-ENABLE beamlines are supported in part by the National Institutes of Health, National Institute of General Medical Sciences, grant P30 GM124169.

## Results

### Identification of a WYL-family transcription factor associated with CBASS systems

To identify novel regulators of anti-phage signaling in CBASS systems, we manually inspected a set of Type III CBASS systems encoded by different *E. coli* isolates. We identified an uncharacterized gene in a CBASS system from *E. coli* strain upec-117, which is predicted to possess both a winged helixturn-helix (wHTH) DNA binding domain and a WYL domain, named for a conserved three-amino acid motif (tryptophan-tyrosine-leucine) in this domain (26). We named this gene *capW* (<u>C</u>BASS-<u>a</u>ssociated <u>p</u>rotein with <u>W</u>YL domain) **(Fig. 1a).** To identify other CBASS systems with *capW*, we searched for homologs of *E. coli* upec-117 *capW* within 10 kb of 6,233 previously-identified CBASS systems in diverse bacteria and archaea (3), and identified 160 CBASS systems encoding this protein **(Table S1).** In these systems, the *capW* gene is consistently found upstream of the core CBASS genes and encoded on the opposite strand **(Fig. 1a).** CapW is associated with all major CBASS types, including Type I (40 systems), Type II (59 systems), and Type III (56 systems) **(Fig. 1b).** These systems encode diverse putative effector proteins, including the phospholipase CapV, endonucleases Cap4 and NucC, and transmembrane proteins Cap14 and Cap15 **(Fig. 1a).** The *E. coli* upec-117 CBASS system encodes three uncharacterized genes: a predicted effector similar to MTA/SAH-family nucleoside phosphorylases (10) that we named Cap17; an uncharacterized predicted 3’-5’ exonuclease that we named Cap18, and a predicted three-transmembrane helix protein that we named Cap19 **(Table S2).**

In other proteins, the WYL domain adopts an SH3 β-barrel fold and has been proposed to function in ligand binding in multiple contexts, including in the regulation of bacterial immunity as part of CRISPR/Cas systems and other immune systems (26,27). This family includes WYL1, a dimeric WYL domain-containing protein that binds single-stranded RNA and positively regulates Cas13d in a Type VID CRISPR-Cas system (28,29). The largest family of bacterial WYL domain-containing proteins possess CapW’s domain structure with an N-terminal wHTH domain, a central WYL domain, and a conserved C-terminal domain termed WCX (WYL C-terminal extension). The structural mechanisms of these proteins, and their roles in bacterial signaling, are largely unknown. The most well-characterized members of this family are PafB and PafC, which together regulate the DNA damage response in mycobacteria (30,31). A recent structure of a naturally-fused PafBC protein from *Arthrobacter aurescens* in the absence of bound ligand or DNA (32) revealed an asymmetric overall structure, leaving unanswered the question of how these proteins’ DNA binding propensity may be regulated by ligand binding. Overall, our bioinformatics data suggest that CapW is a ligand-responsive transcription factor that may regulate expression of its associated CBASS operon in response to phage infection. Moreover, CapW represents a large family of uncharacterized bacterial transcription factors involved in diverse signaling pathways.

### CapW specifically binds the CBASS promoter region

To determine whether CapW is a transcription factor that controls CBASS expression, we first purified the protein and tested its binding to the shared promoter region between the CBASS core genes and CapW. We purified three CapW proteins, from *E. coli* upec-117 (*Ec* CapW), *Stenotrophomonas maltophilia* C11 (*Sm* CapW; 41% identical to *Ec* CapW), and *Pseudomonas aeruginosa* PA17 (*Pa* CapW; 64% identical to *Ec* CapW). All three proteins are associated with Type III CBASS systems **(Fig. 1a, S1a).** By size exclusion chromatography coupled to multi-angle light scattering (SEC-MALS), all three CapW proteins form homodimers in solution **(Fig S1b-d).**

Using an electrophoretic mobility shift assay (EMSA), we found that *Sm* CapW robustly binds to its cognate promoter region **(Fig. 2a).** As *Sm* CapW is a homodimer, we searched its promoter for palindromic sequences that would represent a likely binding site for a two-fold symmetric CapW dimer. We identified two imperfect palindromes within the *S. maltophilia* C11 CBASS promoter region **(Fig. 2b),** and used a fluorescence polarization assay to demonstrate specific binding to a 24-bp sequence that overlaps the promoter’s −10 site **(Fig. 2e).** Next, we searched the promoter regions of *E. coli* upec-117 and *P. aeruginosa* PA17 CBASS for similarly-positioned palindromic sequences. We identified a 21-bp imperfect palindrome in the promoter of *E. coli* upec-117 CBASS, and a 19-bp imperfect palindrome in the promoter of *P. aeruginosa* PA17 CBASS, both of which overlap their promoters’ −10 sites **(Fig. 2c, 2d).** Using fluorescence polarization, we found that *Ec* CapW and *Pa* CapW specifically bind these sequences **(Fig. 2f, 2g).** Based on these data, we conclude that CBASS-associated CapW proteins bind palindromic sequences that overlap the −10 sites within the promoters of their cognate CBASS operons. Because the −10 site represents a key binding site for σ-factors that determine the specificity of RNA polymerase-promoter binding (33), this finding suggests that CapW acts as a transcriptional repressor by inhibiting binding of σ-factors and RNA polymerase to the CBASS promoter.

### Structure and DNA binding mechanism of CapW

We next crystallized and determined the structure of Sm CapW to 1.89 Å resolution **(Table S3).** As expected from our SEC-MALS analysis **(Fig. S1b),** CapW crystallized as a homodimer with each chain possessing an N-terminal wHTH domain, a central WYL domain, and a C-terminal WCX domain **(Fig. 3a).** The CapW homodimer adopts a distinctive domain-swapped overall architecture with the N-terminal wHTH domains adjacent to one another, followed by extended linkers reaching across the dimer such that each protomer’s WYL domain interacts primarily with the wHTH domain of the opposite protomer **(Fig. 3a).** The C-terminal WCX domain adopts an extended ⍰-β fold, and each protomer’s WCX domain reaches back across the top of the dimer to interact with the WYL domain of the opposite protomer **(Fig. 3a).** The two WCX domains define a groove across the top of the CapW dimer that extends between the putative ligand-binding sites of each WYL domain (see next section).

The overall architecture of CapW is equivalent to that of another recently-discovered bacterial defense-associated transcription factor, BrxR. Two parallel studies report the discovery of BrxR as a regulator of BREX anti-phage systems, and determine structures of the protein from two different bacteria (13,14). These structures reveal that BrxR and CapW share a common domain organization and overall domain-swapped architecture, with wHTH domains on one face of the dimer and a putative ligand-binding surface on the opposite face that is made up of WYL and WCX domains.

In DNA-free structures of both CapW and BrxR, the wHTH domains of the two protomers are positioned adjacent to one another, with their DNA-binding surfaces aligned on one face of the dimer. We modelled a DNA-bound structure of CapW based on a structure of *Acinetobacter* BrxR bound to its cognate palindromic DNA sequence (13), revealing that the two wHTH domains in the CapW dimer are perfectly aligned to bind a palindromic DNA sequence ~20 base pairs in length **(Fig. 3b),** close to the length of palindromic sequences we identified as CapW binding sites **(Fig. 2b).**

Based on our model of DNA-bound CapW, we designed two mutants to disrupt DNA binding: a single Arg32 to alanine mutant (*Sm* CapW^R32A^), and a triple mutant with Ser42, Gln45, and Ser47 all mutated to alanine (*Sm* CapW^SQS-AAA^) **(Fig. 3c).** While Arg32 is highly conserved in CapW but not in BrxR, Ser42/Gln45/Ser47 are conserved in BrxR and all three residues are directly involved in DNA binding (13). In *Sm* CapW, both CapW^R32A^ and CapW^SQS-AAA^ eliminated detectable binding of the protein to its binding site in the CBASS promoter **(Fig. 3d).** Together, these data suggest that our structure of Sm CapW represents a DNA-binding competent conformation of CapW.

### The structure of *P. aeruginosa* PA17 CapW reveals a non-DNA binding conformation

The central WYL domain of CapW adopts an Sm-type SH3 β-barrel fold similar to bacterial Hfq (<u>h</u>ost <u>f</u>actor for RNA bacteriophage <u>Q</u>β replication) proteins, which bind small RNAs (34,35). Other WYL domain containing proteins have been shown to bind single-stranded RNA (29) or DNA (36), suggesting that the CapW WYL domain may also bind nucleic acids and/or a small molecule ligand. The WYL domain is named for a set of three highly-conserved amino acids, tryptophan-tyrosine-leucine, located on one of the domain’s β-strands. In CapW, the tryptophan residue is highly conserved, while the typical tyrosine residue in WYL domains is replaced by a highly conserved histidine (His171 in *Sm* CapW; **Fig. S2a-b).** This histidine residue is solvent-exposed on the top face of the CapW dimer, and is surrounded by a cluster of highly-conserved hydrophobic and polar residues including Tyr145, Ser147, Trp156, Arg169, Arg183, Phe185, and Arg189 (Sm CapW residue numbering; **Fig 4a,c, S2a-b, S3).** Comparing these residues to a sequence logo constructed from PFAM13280, which represents over 18,000 WYL-WCX domain proteins, we found that *Sm* CapW Tyr145 aligns with a highly-conserved tyrosine in this family, and Arg183/Phe185/Arg189 are situated in a region that shows high conservation in the broader WYL domain family **(Fig. S2b).** Based on this conservation and on the previously-identified ligand binding role for WYL domains (26,29,36), we propose that this conserved surface on the CapW WYL domain may bind a nucleic acid or small-molecule ligand. Notably, the two putative ligand-binding sites on CapW are situated near one another on the top face of the dimer at either end of a groove defined by the two WCX domains **(Fig. 4a).** The positioning of these sites suggest that they may cooperate to bind an extended nucleic acid ligand.

A major question for the function of CapW and related transcription factors is how these proteins are regulated in order to sense and respond to bacteriophage infection. Based on the putative ligand-binding role for the WYL domain, a compelling model is that ligand binding induces a conformational change that alters the ability of CapW to bind DNA. We determined a crystal structure of a second CapW protein, *Pa* CapW, that sheds light on this potential mode of regulation. We crystallized and determined a 2.3 Å resolution crystal structure of *Pa* CapW in a low-pH condition (pH 5.0). The structure reveals that *Pa* CapW shares the same overall architecture as *Sm* CapW, forming a homodimer of protomers with wHTH, WYL, and WCX domains **(Fig. 4b).** The symmetric arrangement of WYL domains linked by WCX domains is similar to *Sm* CapW, but in *Pa* CapW the WYL domains are each rotated ~10° downward and inward toward the wHTH domains. This motion is accompanied by a slight widening of the groove between the WYL domains’ ligand-binding sites and bordered by the two WCX domains **(Fig. 4b),** and also by a rearrangement of the C-termini of each WCX domain. In *Sm* CapW, the WCX domain C-terminus forms two short ⍰-helices with an intervening loop that folds along the top of the domain **(Fig. 4a).** In *Pa* CapW, this region undergoes a domain swap to fold against the opposite protomer’s WCX domain **(Fig. 4b).** Finally, we observe that a salt bridge between a conserved arginine in the WYL domain and an aspartate on the WCX domain is broken in the *Pa* CapW structure, enabling the WCX domain to move out and away from the dimer-related WYL domain **(Fig. 4d).**

The highly-conserved tryptophan residue of the WYL domain family is positioned on the opposite face of the WYL domain compared to the putative ligand binding site, and in *Sm* CapW this residue (Trp170) packs against an ⍰-helix in the extended wHTH-WYL domain linker (⍰4; **Fig. S4a).** In our structure of *Pa* CapW, the WYL domains pinch inward by ~6 Å **(Fig. 4b),** positioning the equivalent tryptophan residues (Trp181) and the nearby ⍰5 helices too close to one another to accommodate the ⍰4 helices in their original configuration. As a result, the ⍰4 helix of each CapW protomer rotates downward away from the WYL domains **(Fig. S4b).** This change, and the motion of the WYL domains in general, in turn cause a striking ~70° rotation of each wHTH domain compared to its position in *Sm* CapW **(Fig. 4b).** Whereas the wHTH domains are aligned for cooperative DNA binding in our structure of *Sm* CapW, in *Pa* CapW they are completely misaligned and would be unable to bind a contiguous DNA sequence.

Our biochemical data shows that *Pa* CapW binds DNA with an affinity equivalent to that of Sm CapW or *Ec* CapW, yet our structure of this protein reveals a conformation that is clearly unable to bind DNA. We propose that the *Pa* CapW crystal structure reveals a conformational state equivalent to that of ligand-bound CapW, perhaps induced by the low pH of the crystallization condition or by proteinprotein packing interactions. Comparing the DNA-binding and non-DNA binding states of CapW reveals a key role for the WYL domain’s conserved tryptophan residue in driving conformational changes between these two states.

### CapW is a transcriptional repressor for CBASS

Our identification of CapW binding sites overlapping the −10 site of the CBASS promoter suggested that CapW binding may repress CBASS transcription by blocking σ-factor/RNA polymerase binding to this site. To test this idea, we generated an expression reporter system for *Ec* CapW. We first generated DNA-binding mutants of *Ec* CapW equivalent to *Sm* CapW^R32A^ (*Ec* CapW R43A) and CapW^SQS-AAA^ (S53A/Q56A/S58A), and found that both mutants eliminated detectable binding of *Ec* CapW to its palindromic site by fluorescence polarization **(Fig. 5a).** We next constructed an expression reporter system with the *E. coli* upec-117 *capW* gene and CBASS promoter linked to a gene encoding GFP **(Fig. 5b).** To track CapW expression directly in this system, we also fused a C-terminal FLAG tag to the *capW* gene. We measured both GFP and CapW-FLAG expression using Western blots, in the presence and absence of *capW* or a DNA-binding mutant. In the presence of wild-type *capW*, the levels of both GFP and CapW-FLAG were nearly undetectable by Western blotting **(Fig. 5b).** In contrast, disrupting *capW* or eliminating DNA binding through the R43A or SQS-AAA mutants resulted in strong expression of GFP **(Fig. 5b).** We observed a similar increase in expression of CapW-FLAG in both DNA-binding mutants, suggesting that CapW regulates its own transcription in addition to that of the core CBASS genes. We could identify a promoter in the *E. coli* upec-117 CBASS operon that is oriented in the reverse direction compared to the promoter driving core CBASS expression **(Fig. S5).** This promoter likely drives expression of *capW* and the system’s likely effector *cap17*, and our data suggests that CapW binding to the CBASS promoter region affects transcription in both directions. Overall, these data show that CapW is a strong transcriptional repressor, capable of repressing transcription bidirectionally from the CBASS promoter.

Our data suggest that in its unliganded state, CapW strongly represses CBASS transcription by binding the operon’s promoter. To test whether this repression is released upon phage infection, potentially by ligand binding to the protein’s WYL domain, we used our GFP reporter system to measure expression after infection with phage λ. With wild-type *capW*, GFP expression was undetectable in uninfected cells, but was detectable within 30 minutes of phage λ infection, and increased through 120 minutes post-infection **(Fig. 5d).** We generated a series of point mutants to five polar and charged residues in the putative ligand-binding site of *Ec* CapW (based on the structure of *Pa* CapW; **Fig. 5c),** and found that all of these mutants showed either no detectable expression increase after infection, or extremely low/delayed expression compared to the system encoding wild-type CapW **(Fig. 5d).** The equivalent point mutants of Sm CapW show no loss of DNA binding affinity in vitro **(Fig. S3e),** supporting a model in which these mutants render CapW a constitutive transcriptional repressor. These data reveal a key role for the WYL domain’s putative ligand binding site in CapW-dependent control of CBASS expression.

We next tested whether a CapW-mediated CBASS expression increase is required for phage protection by the *E. coli* upec-117 CBASS system. We found that this system shows strong protection against infection by a strain of phage λ that obligately undergoes the lytic infection cycle (λ cI-) **(Fig. 5e)** (37). This protection depends on the system’s cGAS-like enzyme CdnC and its putative effector, the predicted MTA/SAH-family nucleoside phosphorylase Cap17 **(Fig. 5e).** We found that a catalytic-dead mutant of this system’s predicted 3’-5’-exonuclease (Cap18) did not affect phage protection, but that introduction of a stop codon into the gene encoding the uncharacterized transmembrane protein (Cap19) strongly affected phage protection **(Fig. 5e).** Surprisingly, we found that the CapW WYL domain mutants that constitutively repress CBASS expression do not compromise the system’s ability to protect against phage λ **(Fig. 5e).** Thus, the low levels of CBASS expression present in uninfected cells (and in CapW WYL-domain mutants) are sufficient for a robust anti-phage response.

Despite repeated attempts, we were unable to generate mutant *E. coli* upec-117 CBASS constructs that either disrupted *capW* or eliminated its DNA binding activity. After mutagenesis, clones containing these mutations consistently also showed large deletions of critical regions of the CBASS operon (not shown). Our inability to isolate these mutants in the context of a full CBASS system, when the same mutants are readily obtainable in our GFP reporter system, suggests that elimination of CapW-mediated repression and the resulting high-level expression of CBASS genes is likely toxic to host cells. This model is consistent with a parallel study on the related BrxR transcription factor, in which deletion of BrxR leads to increased expression of BREX genes and toxicity to host cells (13,14).

## Discussion

Bacterial CBASS immune systems are highly diverse, and an emerging theme in these systems is that they encode multiple redundant regulatory mechanisms in order to, for example, prevent antiphage signaling and associated cell killing outside the context of infection. All CBASS systems encode a cGAS-like oligonucleotide cyclase that likely possesses an inherent phage-dependent activation mechanism (4,10). Type II, III, and IV CBASS systems additionally encode regulators - many of which represent ancestral forms of important eukaryotic signaling proteins - that provide a second level of control over the activity of their cognate cGAS-like enzymes (7,10). Here, we identify a third mode of regulation in some CBASS systems: a WYL domain-containing transcription factor, CapW. We show that CapW strongly represses CBASS expression by binding the CBASS promoter region, and that repression can be released upon phage infection in a manner that depends on the putative ligand-binding site in the protein’s WYL domain. Our two structures suggest that ligand binding to the protein’s WYL domain causes a conformational change in the WYL and WCX domains that triggers a large-scale rotation of the wHTH domains to release the protein from DNA **(Fig. 6).**

**Figure 6.**
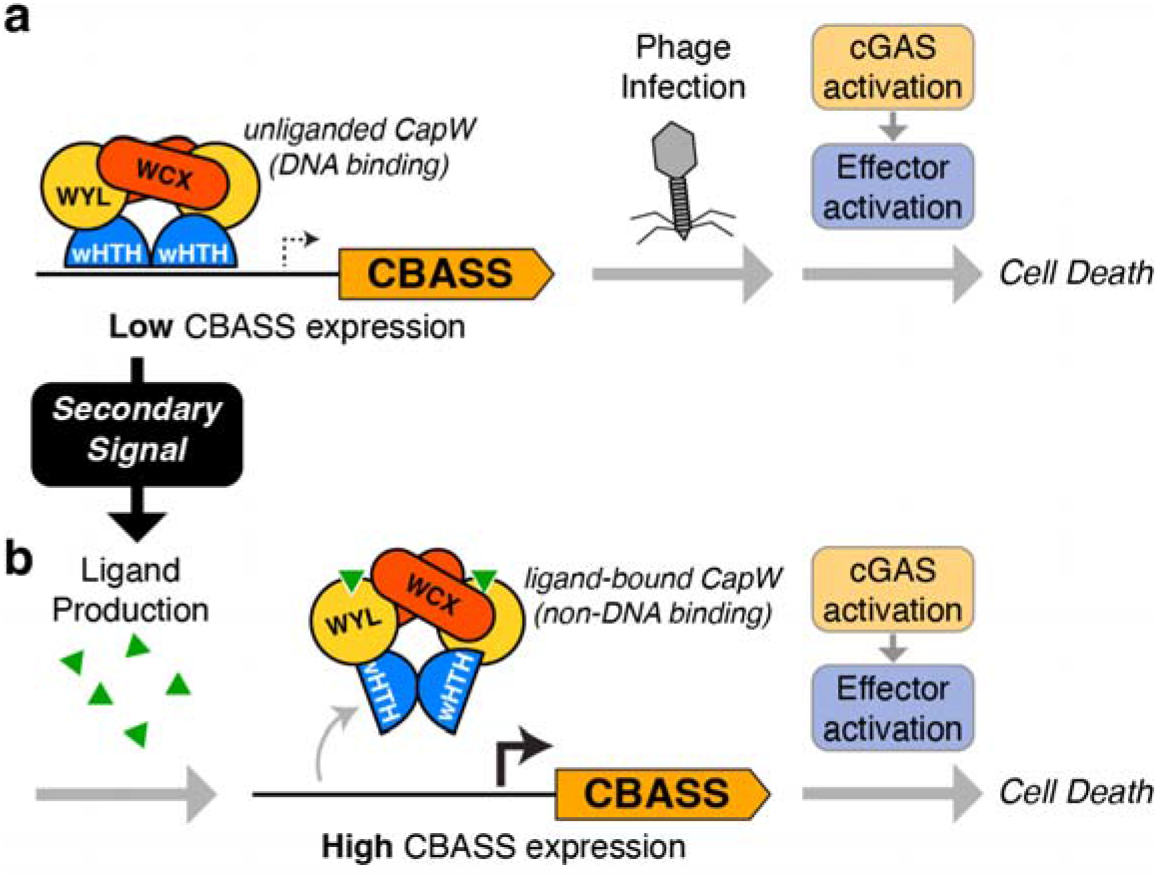
Model for CapW function in CBASS. (a) In an uninfected cell, the dimeric CapW transcription factor binds the promoter of its cognate CBASS system and maintains its expression at a low level. Upon phage infection, cGAS is activated to produce a second messenger signal that in turn activates the system’s effector, killing the infected cell. (b) In response to a secondary stress signal that produces a small-molecule or nucleic acid ligand, CapW binds the ligand and undergoes a conformational change to release it from DNA. The resulting loss of transcriptional repression causes high-level CBASS expression and associated cell death even in the absence of the system’s primary phage infection signal.

Mutation of conserved residues in CapW’s WYL domain eliminate CapW-dependent CBASS derepression upon phage λ infection, yet paradoxically do not affect the anti-phage activity of CBASS. This observation, combined with our inability to delete copW in the *E. coli* upec-117 CBASS system, suggests a two-pronged model for CBASS action in CapW-containing systems. We propose that the system’s primary mode of action responds directly to phage infection, through either cGAS’s inherent phage-sensing activity or the action of known regulators, and kills the host cell to prevent phage replication **(Fig. 6a).** This mode does not require high-level expression of CBASS proteins, as demonstrated by the robust anti-phage activity of systems containing CapW mutants lacking the ability to respond to phage infection **(Fig. 5d-e).** In CBASS systems encoding CapW, we propose a secondary mode in which an unknown stress signal causes production of a CapW-binding ligand, releasing CapW from DNA to drive high-level expression of CBASS genes **(Fig. 6b).** Since our data and prior studies suggest that high-level CBASS expression is inherently toxic to host cells (38), this secondary mode would also result in cell death even in the absence of the primary phage trigger(s) sensed by the system. Thus, CapW may enable a single CBASS system to respond directly to phage infection (through its primary mode of action) and also respond to other, as-yet unidentified stress signals. Phage λ infection does trigger CapW-mediated transcription de-repression **(Fig. 5d),** with the strongest effect observed around the time host cells undergo phage-mediated lysis (39). An important future direction will be to isolate the molecular signals that mediate this de-repression, providing insight into the stress pathways that act on CapW’s secondary signaling mode.

CapW shares a similar overall architecture to another recently-identified transcription factor, BrxR, which controls the expression of BREX immune systems (13,14). Like CapW, BrxR binds BREX promoter sequences to repress expression in uninfected cells (13,14). Curiously, introduction of an early stop codon into *Adnetobacter* BrxR to eliminate protein production does not compromise the anti-phage activity of its cognate BREX system (13). Based on this observation, Luyten et al. suggest that BrxR may activate BREX as a “second line of defense” by responding to a ligand produced by a stress pathway or product of other defense systems like CRISPR/Cas or restriction-modification (13). We propose that the allosteric mechanism we identify for ligand-induced conformational changes in CapW also applies to BrxR, and to the larger family of bacterial defense-associated WYL domain transcription factors. CapW/BrxR-like transcription factors are associated with a variety of bacterial immune systems including CRISPR-Cas and restriction-modification systems (14), and all of these proteins share a conserved ligandbinding surface **(Fig. S2c),** suggesting that they may bind the same or similar ligands. Moreover, these proteins all share the conserved tryptophan residue after which the WYL domain is named, which we implicate in allosteric communication between WYL/WCX and wHTH domains in CapW.

Outside bacterial immune systems, the WYL domain and WYL-WCX domain pair likely play a range of signaling roles. For example, the transcription factor PafBC possesses a tandem array of wHTH, WYL, and WCX domains, and plays a role in regulating the response to DNA damage in mycobacteria (32). The Cas13d regulator WYL1 shares a similar domain structure with an N-terminal ribbon-helix-helix domain, a central WYL domain, and a C-terminal dimerization domain (28,29). Examining sequence conservation in PFAM13280, which includes over 18,000 bacterial proteins that possess the WYL-WCX domain pair embedded in a variety of protein scaffolds, reveals that the ligand-binding site is much more variable across this family than within the smaller group of proteins associated with immune systems **(Fig. S2b).** Thus, while immune system-associated WYL proteins likely bind a common ligand, other family members have likely evolved to bind a variety of ligands to control diverse signaling pathways. Key directions for future work will be to determine the range of ligands bound by diverse WYL proteins, how ligand binding is coupled to conformational changes within different protein scaffolds, and how these conformational changes are coupled to signaling.

## Data Availability

The datasets produced in this study are available in the following databases:

Protein structure data: Protein Data Bank IDs 7TB5 (http://dx.doi.org/10.2210/pdb7tb5/pdb) and 7TB6 (http://dx.doi.org/10.2210/pdb7tb6/pdb).

X-ray diffraction data: SBGrid Data Bank IDs 869 (https://data.sbgrid.org/dataset/869/) and 870 (https://data.sbgrid.org/dataset/870/).

## Acknowledgements

We thank A. Whiteley for the gift of a plasmid encoding the *E. coli* upec-117 CBASS system, and members of the Corbett lab for helpful discussions and comments on the manuscript.

## Supplemental Data

**Figure S1.**
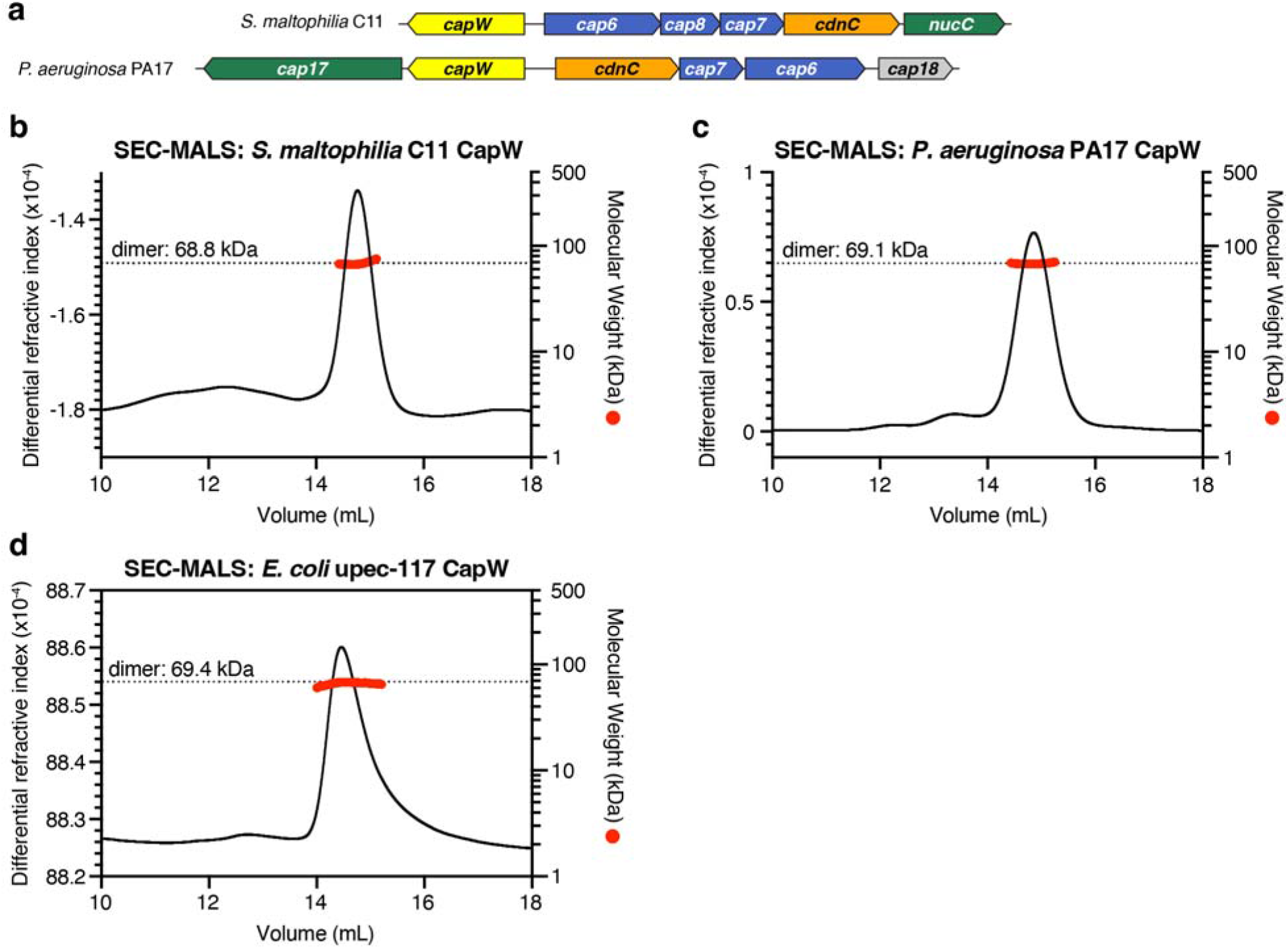
Identification and characterization of CBASS-associated CapW. (a) Operon schematics of CBASS systems from *P. aeruginosa* PA17 and S. *maltophilia* C11, colored as in **Fig. 1a.** (b) Size exclusion chromatography coupled to multi-angle light scattering (SEC-MALS) analysis of purified *Sm* CapW. The measured molecular weight (62.2 kDa; red line) is consistent with a homodimer (68.8 kDa; dotted black line). (c) SEC-MALS analysis of purified *Pa* CapW. The measured molecular weight (68.2 kDa; red line) is consistent with a homodimer (69.1 kDa; dotted black line). (d) SEC-MALS analysis of purified *Ec* CapW. The measured molecular weight (66.1 kDa; red line) is consistent with a homodimer (69.4 kDa; dotted black line).

**Figure S2.**
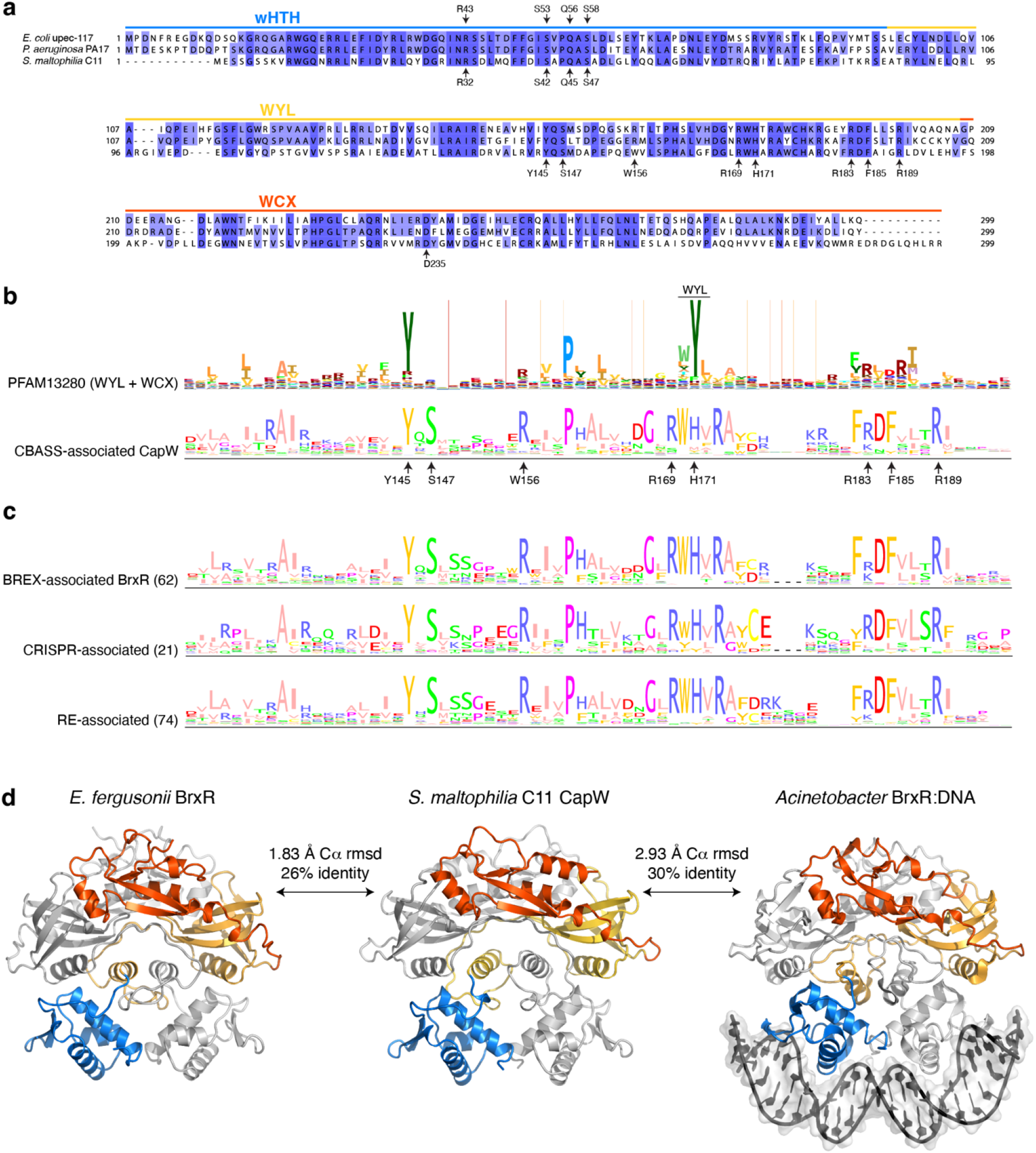
Sequence alignments of CapW and related transcription factors. (a) Sequence alignment of CapW from *E. coli* upec-117, *P. aeruginosa* PA17, and *S. maltophilia* C11. Blue, yellow, and orange lines indicate the extent of the wHTH, WYL, and WCX domains, respectively. wHTH domain residues involved in DNA binding are labeled for *Ec* CapW and *Sm* CapW. Also labeled are putative ligand-binding and allosteric-regulation residues in the WYL and WCX domains of Sm CapW. (b) Sequence logos of PFAM13280 (assembled from ~18,000 WYL-WCX domain containing proteins; www.pfam.org) and 160 CBASS-associated CapW proteins (generated with JALVIEW from the proteins listed in **Table S1)** (41). Key putative ligand-binding site residues of Sm CapW are labeled. (c) Sequence logos of BrxR (62 sequences) and related WYL domain transcription factors associated with CRISPR/Cas (21 sequences) and restriction-modification systems (74 sequences), as identified in Picton et al. (14). (d) Comparison of the Sm CapW structure (center) with those of *E.fergusonii* BrxR (PDB ID 7QFZ; (14)) and DNA-bound *Acinetobacter* BrxR (PDB ID 7T8K; (13)). All three proteins are shown in the same orientation, with one protomer gray and the other colored as in **Fig. 3a.** Despite relatively low sequence identity with BrxR, *Sm* CapW shows high overall structural similarity to both structures.

**Figure S3.**
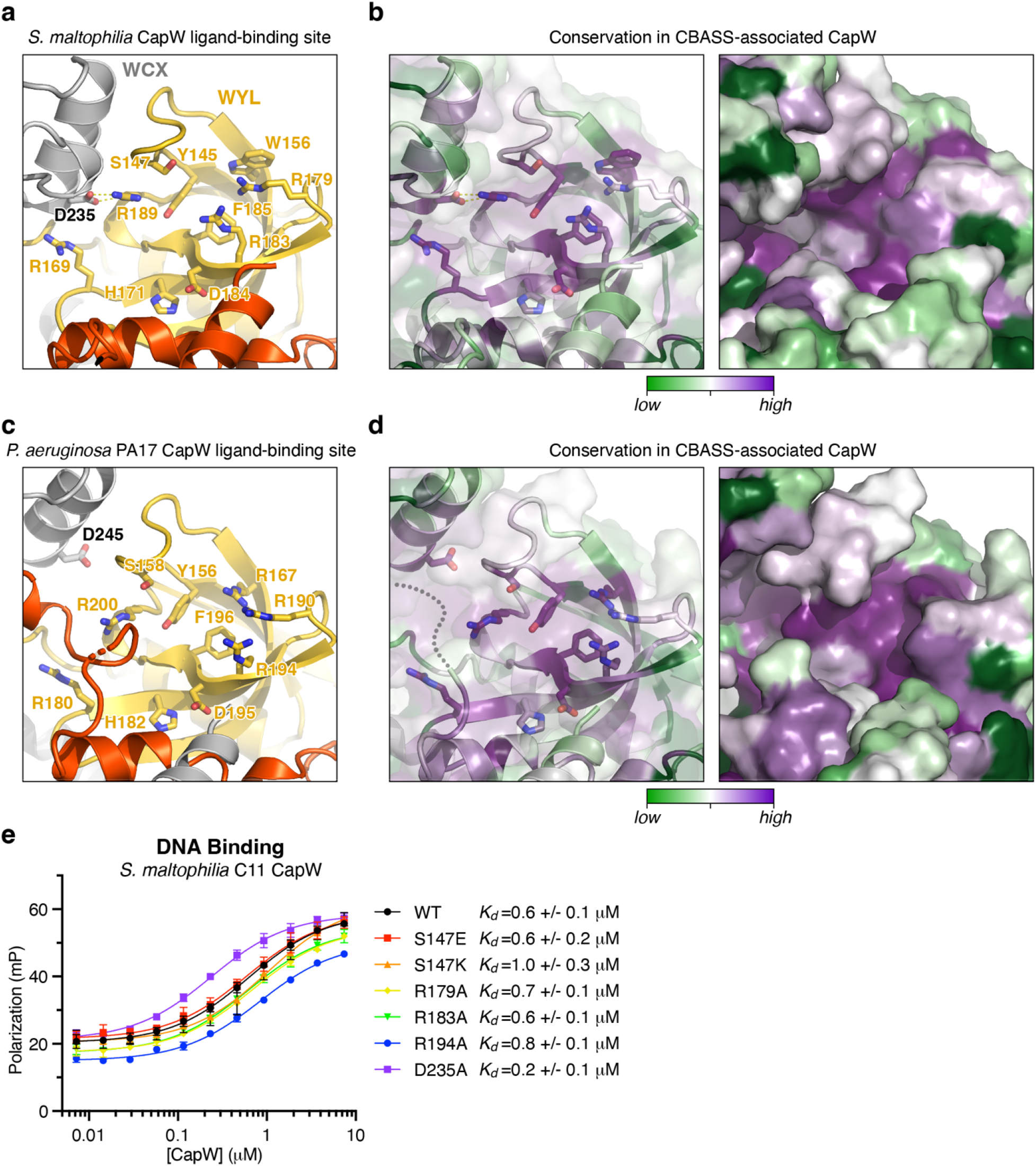
Surface conservation of the CapW putative ligand binding site. (a) Top-down view of the putative ligand-binding site of *Sm* CapW, with WYL domain colored yellow and dimer-related WCX domain gray. (b) View equivalent to panel (a), colored by conservation among 160 CBASS-associated CapW proteins (green: poorly conserved; purple: highly conserved); conservation scores calculated using the ConSurf server (42). (c) Top-down view of the putative ligand-binding site of *Pa* CapW, with WYL domain colored yellow and dimer-related WCX domain gray. (d) View equivalent to panel (c), colored by conservation among 160 CBASS-associated CapW proteins. For clarity, residues 276-285 have been removed from these panels (gray dotted line). (e) Fluorescence polarization DNA binding assay with S. *maltophilia* C11 CapW (wild-type and indicated mutants of the conserved putative ligand-binding surface) binding DNA Probe #1 **(Fig. 2b).**

**Figure S4.**
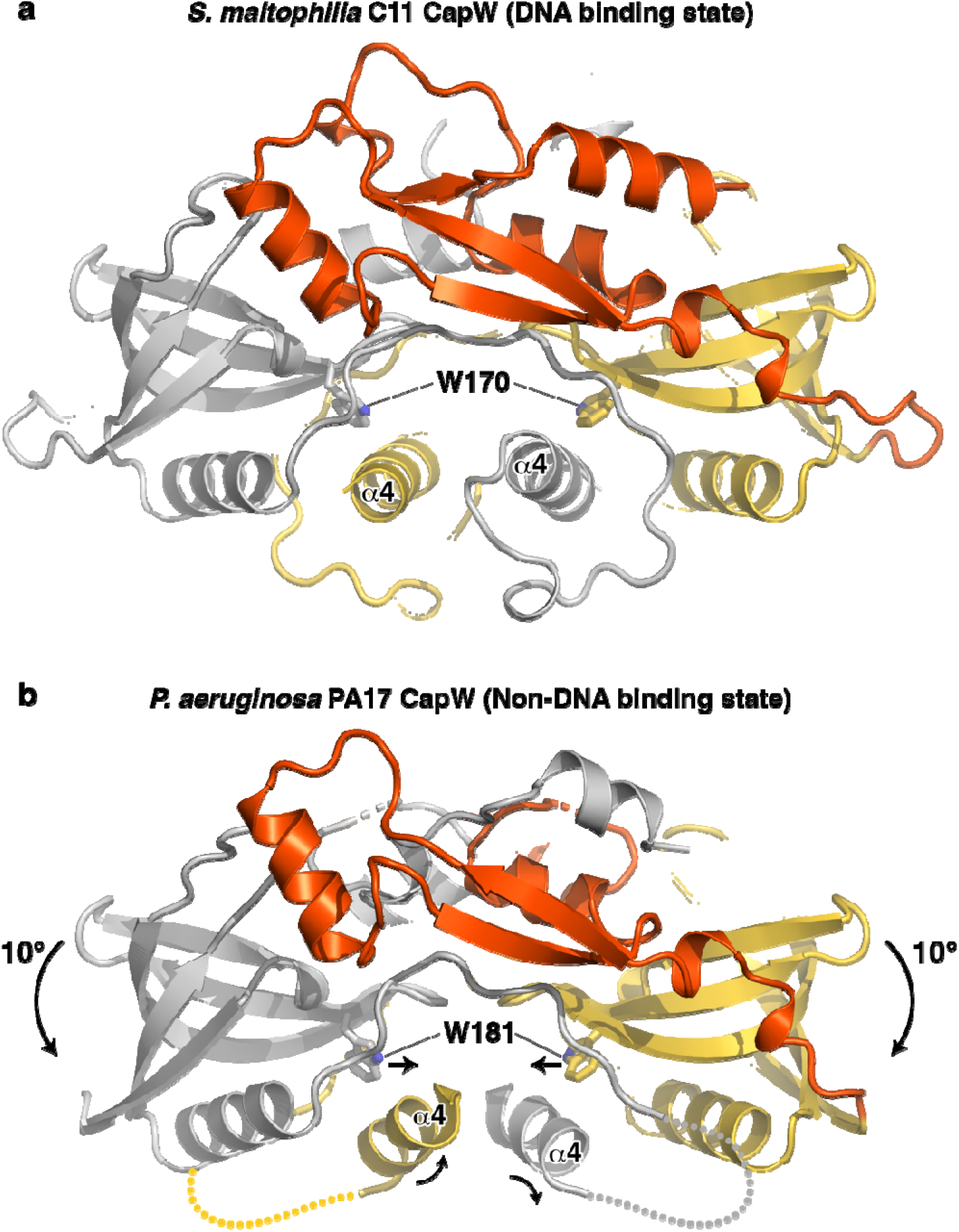
Conformational changes in CapW are driven by WYL domain motions. (a) Front view of the *Sm* CapW dimer, with wHTH domains omitted for clarity. Shown in sticks is residue W170 for both monomers, which packs against the wHTH-WYL linker ⍰4 helix, (b) Front view of the *Pa* CapW dimer, with wHTH domains omitted for clarity. Shown in sticks is residue W181 for both monomers. Compared to the structure of Sm CapW, the two W181 residues are closer together, driving rotation of the ⍰4 helix (indicated by arrows) and rotation of the associated wHTH domains (not shown).

**Figure S5.**
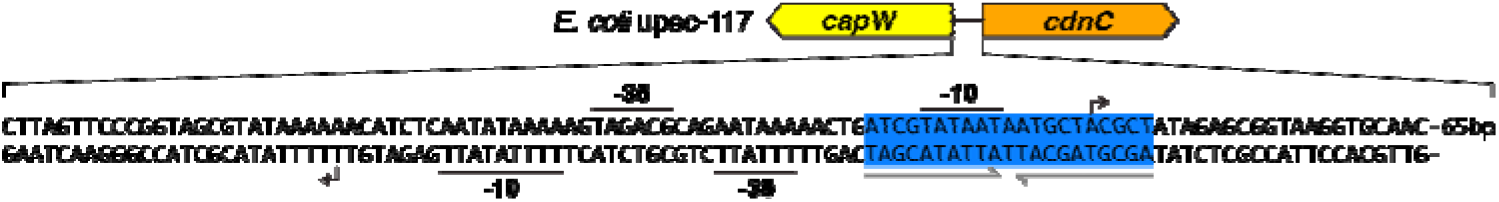
Promoter structure of *E. coli* upec-117 CBASS. Diagram of the *E. coli* upec-117 CBASS promoter showing key sequences (−35 site, −10 site, and transcription start site indicated by arrow) for the top strand (driving core CBASS expression) and the bottom strand (driving *capW* and *cap17* expression). Promoter sequences (−35, −10, and TSS sites) were predicted with the BPROM server (40). Shown in blue is the palindromic sequence that CapW binds.

**Figure S6.**
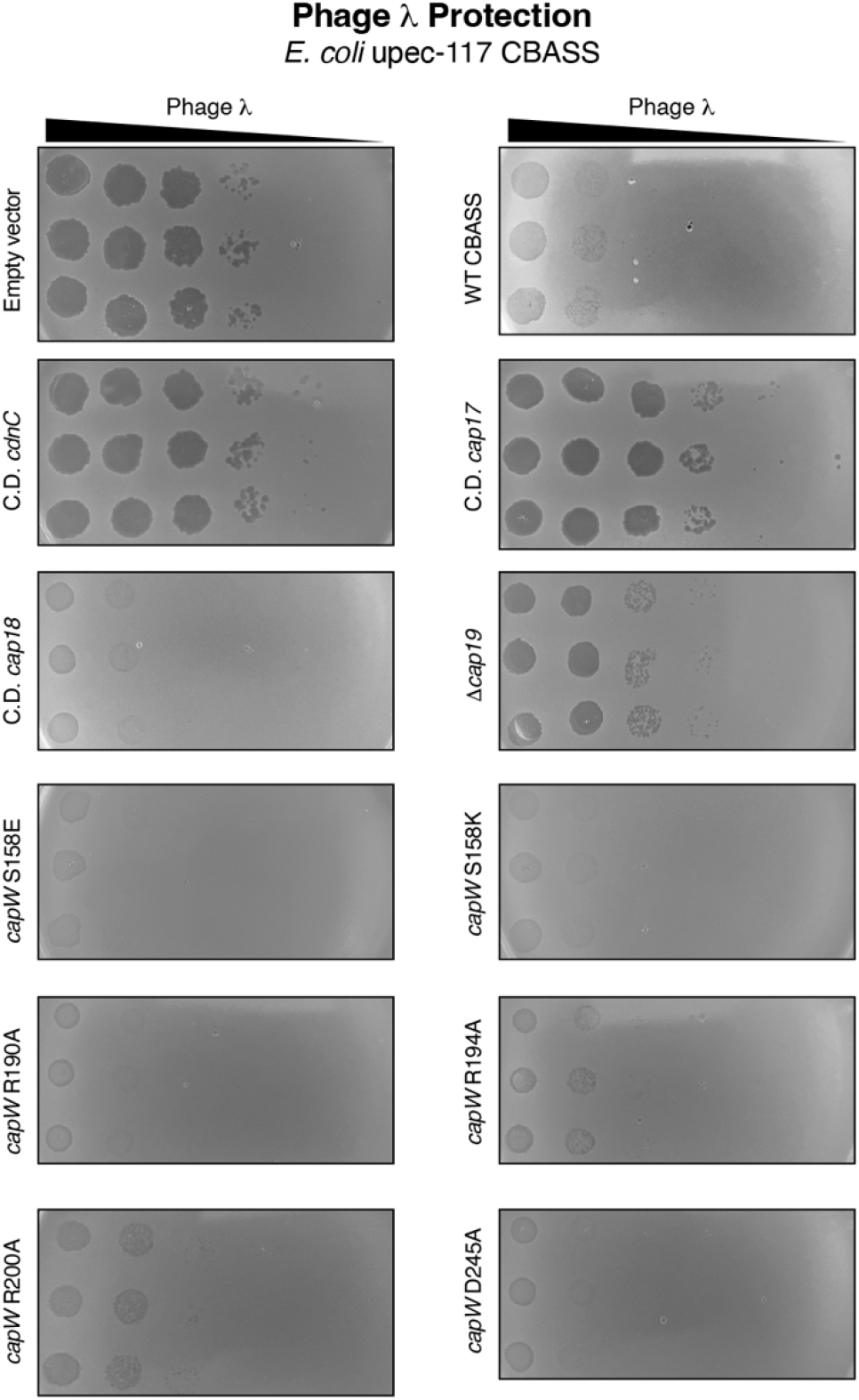
Phage protection by *E. coli* upec-117 CBASS. Plaque assays performed with phage λ cI-on *E. coli* JP313 cells with a vector encoding the *E. coli* upec-117 CBASS system (WT CBASS) or the indicated mutants. Each assay was performed in triplicate with six 10-fold dilutions of phage, then plaques were counted in the highest dilution with visible plaques for each strain. For several strains (C.D. *cap18* and *copW* S158E/S158K/R190A/D245A), plaques uniform clearing was observed for the two most concentrated phage dilutions, but not after. For quantitation **(Fig. 5e),** these were counted as a single plaque in the third dilution.

## Supplemental Tables

**Table S1. CBASS systems with associated CapW**

(*see attached Excel sheet*)

**Table S2.** CBASS-associated regulators and effectors.

| Gene name | Effector/Regulator | Description | Structure(s) | Reference |
| --- | --- | --- | --- | --- |
| Cap1/CapV | Effector | Phospholipase | n/a | (38) |
| Cap2 | Regulator (Type II) | Ubiquitin E2/E1-like | n/a | (4,43) |
| <sup>a</sup> Cap2b | Regulator (Type II - short) | Ubiquitin E2-like | n/a | (4,10) |
| Cap3 | Regulator (Type II) | JAB/JAMM (iso)peptidase-like | n/a | (4,43) |
| Cap4 | Effector | SAVED endonuclease | 6WAN, 6VM6, 6VM5, 6WAM | (44) |
| Cap5 | Effector | HNH endonuclease | 7RWM, 7RWK | (44,45) |
| Cap6 | Regulator (Type III) | Trip13 | 6PB3, 6P8V | (7) |
| Cap7 | Regulator (Type III) | HORMA1 (in 1-HORMA systems); HORMA2 (in 2-HORMA systems) | 6P8O, 6P8P, 6P8R, 6P8S, 6P8U, 6P8Q, 6U7B, 6P8V | (7) |
| Cap8 | Regulator (Type III) | HORMA3 (2<sup>nd</sup> HORMA in 2-HORMA systems) | 6P8S | (7) |
| Cap9 | Regulator (Type IV) | QueC | n/a | (10) |
| Cap10 | Regulator (Type IV) | TGT | n/a | (10) |
| Cap11 | Regulator (Type IV) | N-glycosylase/DNA lyase (OGG) | n/a | (10) |
| Cap12 | Effector | TIR-STING | 6WT4, 6WT5 | (11) |
| Cap13 | Effector | TM-STING | n/a | (11) |
| Cap14 | Effector | TM-SAVED | n/a | (46) |
| Cap15 | Effector | TM-β-barrel | 7N34, 7N35 | (46) |
| Cap16 | Effector | TM-NUDIX | n/a | (46) |
| NucC | Effector | endonuclease | 6P7O, 6P7Q, 6P7P, 6Q1H, 6UXF, 6UXG | (39) |
| <sup>a</sup> Cap17 | Effector | MTA/SAH nucleoside phosphorylase | n/a | (10), this study |
| <sup>a</sup> Cap18 | *unknown* | 3'-5' exonuclease | n/a | (10), this study |
| <sup>a</sup> Cap19 | *unknown* | 3-transmembrane protein | n/a | this study |

<sup>a</sup> Gene names coined in this study.

**Table S3.**
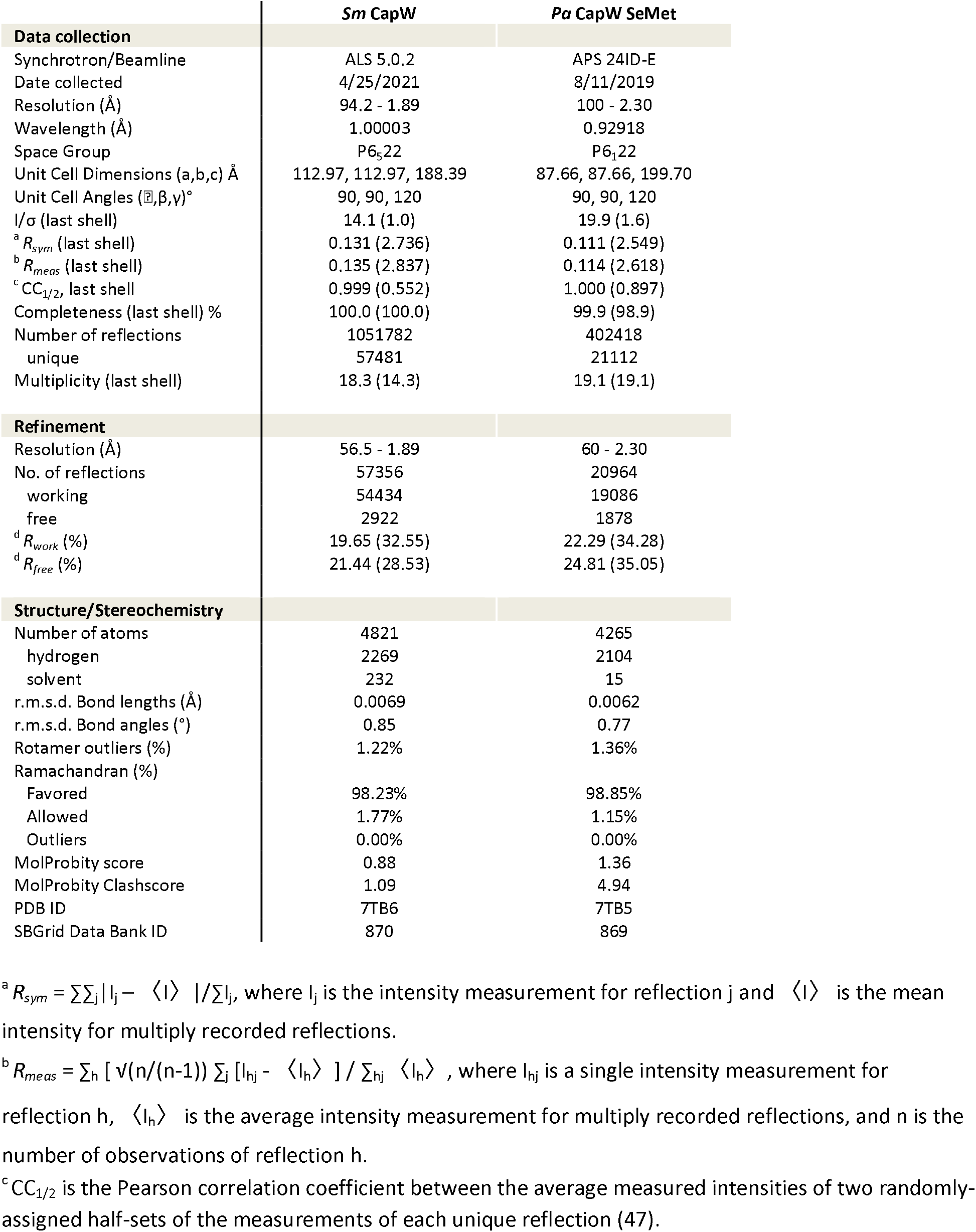
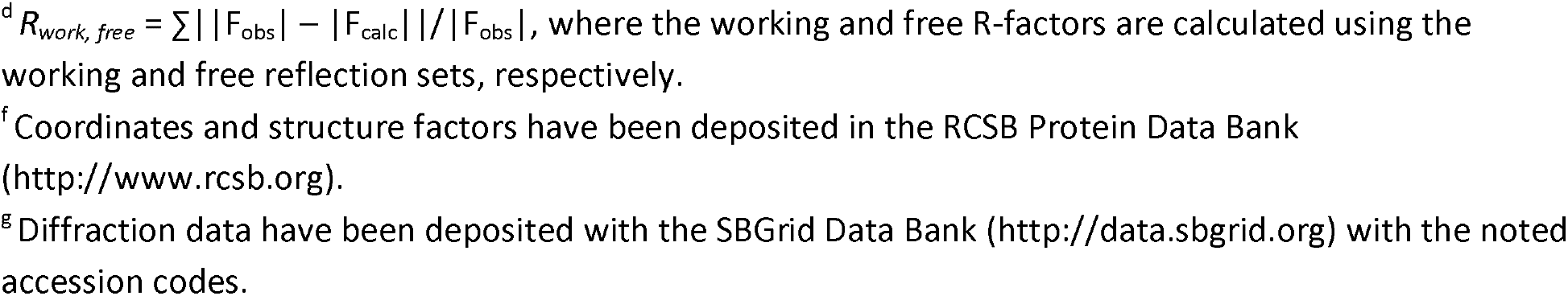
Crystallographic data collection and refinement.

